# Frontal hip exoskeleton assistance does not appear promising for reducing the metabolic cost of walking: A preliminary experimental study

**DOI:** 10.1101/2023.08.22.554212

**Authors:** Jinsoo Kim, Michael Raitor, C. Karen Liu, Steven H. Collins

## Abstract

**Background:** During walking, humans exert a substantial hip abduction moment to maintain balance and prevent pelvic drop. This significant torque requirement suggests that assisting the frontal hip muscles could be a promising strategy to reduce the energy expenditure associated with walking. A previous musculoskeletal simulation study also predicted that providing hip abduction assistance through an exoskeleton could potentially result in a large reduction in whole-body metabolic rate. However, to date, no study has experimentally assessed the metabolic cost of walking with frontal hip assistance.

**Methods:** In this case study involving a single subject (N = 1), a tethered hip exoskeleton emulator was used to assess the feasibility of reducing metabolic expenditure through frontal-plane hip assistance. Human-in-the-loop optimization was conducted separately under torque and position control to determine energetically optimal assistance parameters for each control scheme.

**Results:** The optimized profiles in both control schemes did not reduce metabolic rate compared to walking with assistance turned off. The optimal peak torque magnitude was found to be close to zero, suggesting that any hip abduction torque would increase metabolic rate. Both bio-inspired and simulation-inspired profiles substantially increased metabolic cost.

**Conclusion:** Frontal hip assistance does not appear to be promising in reducing the metabolic rate of walking. This could be attributed to the need for maintaining balance, as humans may refrain from relaxing certain muscles as a precaution against unexpected disturbances during walking. An investigation of different control architectures is needed to determine if frontal-plane hip assistance can yield successful results.

## Background

Hip abductor and adductor muscles play a crucial role in maintaining balance during walking. They are responsible for ensuring proper foot placement after the swing phase [1], which is essential for mediolateral stability [2]. Additionally, these muscles prevent pelvic drop during the single-limb support phase. When the contralateral limb is lifted off the ground, the weight of the upper body creates a significant external adduction moment around the hip joint. To counteract this gravitational moment and stabilize the pelvis relative to the femur, the hip abductors are engaged [3,4]. The magnitude of this biological hip abduction moment is substantial, with peak torque reaching approximately 1.3 Nm/kg [5]. This exceeds the peak sagittal-plane torques of the hip and knee and is comparable to peak ankle plantarflexion torque. This suggests that frontal hip motion could be a promising target for exoskeleton assistance, with the potential to reduce the energetic cost associated with walking.

Musculoskeletal simulations have also indicated the promise of assisting hip abduction for metabolic reduction, but a gap remains between experiment and simulation. For instance, a simulation study by Dembia et al. predicted that a hip abduction exoskeleton could achieve approximately a 13% reduction in whole-body metabolic rate [6]. Interestingly, this reduction ranked as the second largest among the seven different lower limb joint motions examined (i.e., hip flexion/extension, knee flexion/extension, ankle dorsiflexion/plantarflexion, and hip abduction). Despite these encouraging simulation findings, the majority of exoskeletons developed to date have primarily focused on providing assistance in the sagittal plane, neglecting frontal hip assistance [7–15]. Notably, no single study has experimentally demonstrated a metabolic reduction with frontal hip assistance [16,17]. While four exoskeletons have been identified that actively provide frontal hip assistance [18–21], their impact on energy expenditure remains unreported. This highlights the need for further research and exploration of exoskeleton controls and designs that target frontal hip assistance in order to fully exploit their potential benefits in reducing the energetic cost of walking.

Determining the optimal control scheme for frontal hip assistance presents a challenge due to its unique gait function and distinct joint dynamics. Previous exoskeletons have predominantly employed torque control to reduce metabolic cost, prioritizing assistance to the hips and/or ankles in the sagittal plane, as these joints contribute the most positive power during walking [22]. Torque control has proven effective in augmenting hip and ankle power tasks, leading to net metabolic reductions in sagittal plane exoskeleton studies [8–14,23,24]. However, hip abduction/adduction serves a different role in gait, primarily focused on maintaining postural stability and preventing pelvic drop rather than providing substantial positive power to the joint. The focus of frontal plane hip function on posture and support indicates closed-loop position control may also be suitable for delivering frontal plane hip assistance, as it enables precise tracking of the pelvis position relative to the femur. Given the past success of torque control and the functional suitability of position control, both control strategies are promising for frontal plane hip assistance.

Human-in-the-loop optimization (HILO) is a valuable tool for determining exoskeleton control parameters that significantly reduce metabolic cost [8–12,24,25]. While bioinspiration, simulation models, and user feedback-based hand tuning can provide initial suggestions for control parameters, previous research has consistently demonstrated that HILO outperforms these approaches for easily quantifiable outcomes, such as metabolic cost and walking speed [8,11,26]. Selecting effective control parameters for exoskeletons is a complex task due to the difficulty in modeling or predicting human responses [27], the variability in individual responses to exoskeleton assistance [28,29], and the need for extensive training to use exoskeletons effectively [30]. When developing a new exoskeleton or addressing a new task, such as walking with frontal hip assistance, HILO becomes particularly valuable as it can identify effective control parameters that may not be known in advance. Moreover, HILO can enable better comparisons across devices or control approaches because the best versions in each scenario can be found and compared.

The primary objective of this study was to assess the potential impact of frontal hip assistance on reducing the metabolic cost of walking. Additionally, we aimed to compare the effectiveness of torque control and position control in providing frontal hip assistance. We used human-in-the-loop optimization to identify the most effective control parameters for each scheme and evaluated their effects on metabolic rate. During this exploratory phase, we conducted preliminary single-subject tests to gauge the potential promise of this approach. We anticipated that the results would offer insights into the viability of frontal plane assistance and guide the selection of an appropriate controller scheme for tasks that do not require substantial positive power.

## Methods

We used human-in-the-loop optimization to determine frontal hip assistance that minimizes the metabolic cost of walking. The participant wore a hip exoskeleton that controlled hip abduction/adduction [31] and walked on a split-belt treadmill at a speed of 1.25 m/s (Fig. 1a). We tested assistance separately for the torque control and position control schemes. Within each scheme, we optimized the assistance profile and evaluated its effect on metabolic rate. We compared walking with optimized assistance to walking without exoskeleton assistance, as well as natural walking without the exoskeleton, while also evaluating the effects of bio-inspired and simulation-inspired assistance. Finally, we compared the performance of the two control schemes.

**Fig. 1.**
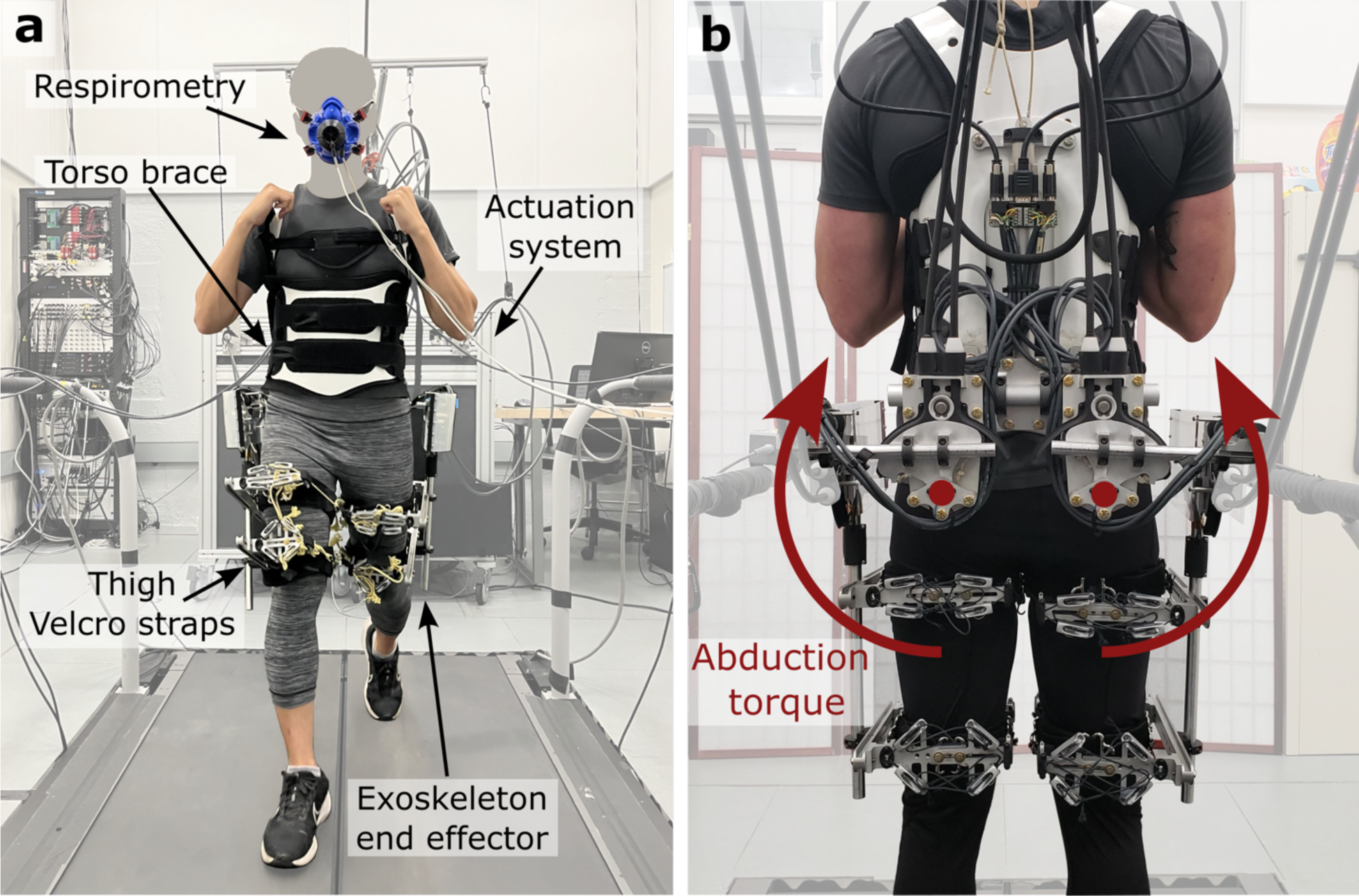
**a,** Experimental setup. A participant walks on a split-belt treadmill wearing a tethered bilateral hip exoskeleton and a respirometry mask. **b,** Rear view of the hip exoskeleton end effector. The exoskeleton is actuated by off-board motors via a Bowden cable transmission.

### Participants

One healthy, young adult (32 years old, male, 168 cm, 60 kg) with prior experience using the device participated in this case study. With prior device experience, the participant was expected to quickly acclimate to both new assistance profiles and control schemes.

### Exoskeleton Hardware

Frontal hip assistance was implemented using a hip exoskeleton emulator (Fig. 1a) [31]. The emulator consisted of an off-board actuation unit equipped with eight motors (Humotech, PA, USA), a hip exoskeleton end-effector, and flexible Bowden cable transmissions connecting these components. The end-effector was attached to the user’s torso using a commercially available torso brace, while each thigh was secured to it via two hook-and-loop Velcro straps. The hip exoskeleton allowed for two degrees of freedom in abduction/adduction and flexion/extension for each leg, resulting in a total of four degrees of freedom for both legs combined. Active assistance was provided for abduction/adduction in both torque and position control schemes (Fig. 1b). In contrast, for flexion/extension, the Bowden cable transmission remained slack (i.e., no resistance), allowing the user to move their limbs freely in the sagittal plane without encountering any applied torque. Due to the lateral placement of thigh attachment rods in the hardware design, which slightly impeded the natural arm swing, the participant grasped the shoulder straps of the torso brace while walking (Fig. 1a). The device had a peak torque capacity of 75 Nm and a total mass of 12.0 kg, including Bowden cables.

### Exoskeleton Control and Control Parameterization

#### Percent Stride Estimation

Heel strike and toe-off were detected by measuring ground reaction forces and moments on a split-belt instrumented treadmill (Bertec, OH, USA). To address the issue of crossovers during frontal hip assistance, we used a combination of vertical ground reaction forces (GRF) and a weighted center of pressure (CoP) in the fore-aft direction. For normal steps, where the left and right heels contacted their respective treadmill belts, our algorithm used simple thresholding on the vertical ground reaction force to detect heel-strike and toe-off events. For crossover steps, the algorithm identified the local minimum (or maximum) in the weighted fore-aft center of pressure during the single (or double) stance phase and classified it as a heel strike (or toe-off), in a method similar to that of Verkerke et al. [32]. The algorithm continuously monitored these two conditions – simple thresholding on vertical GRF or identification of local extrema in fore-aft CoP – without prior knowledge of whether the current step was normal or a crossover, and it activated one of them if the specific criterion was met. This combined approach ensured the timely identification of heel strike and toe-off during normal steps while reliably detecting these gait events even in the presence of crossovers. Subsequently, the algorithm calculated the stride time as the duration between consecutive heel strikes from the same leg and estimated the percent stride based on the time since the last heel strike and the average stride time from the five most recent steps.

#### Torque Control

In the torque control scheme, the exoskeleton was controlled by applying frontal hip joint torques as a function of the percent stride. This torque profile was represented by a piecewise cubic Hermite interpolating polynomial spline with four nodes (Fig. 2a): the onset, the first peak, the second peak, and the offset. Four parameters were used to determine the magnitudes and timings of the two middle nodes, while the timings of the onset and offset nodes were fixed at the heel strike and toe-off, respectively. The minimum and maximum limits of these parameters were established based on user comfort during pilot tests (Additional file 1: Table A1).

**Fig. 2.**
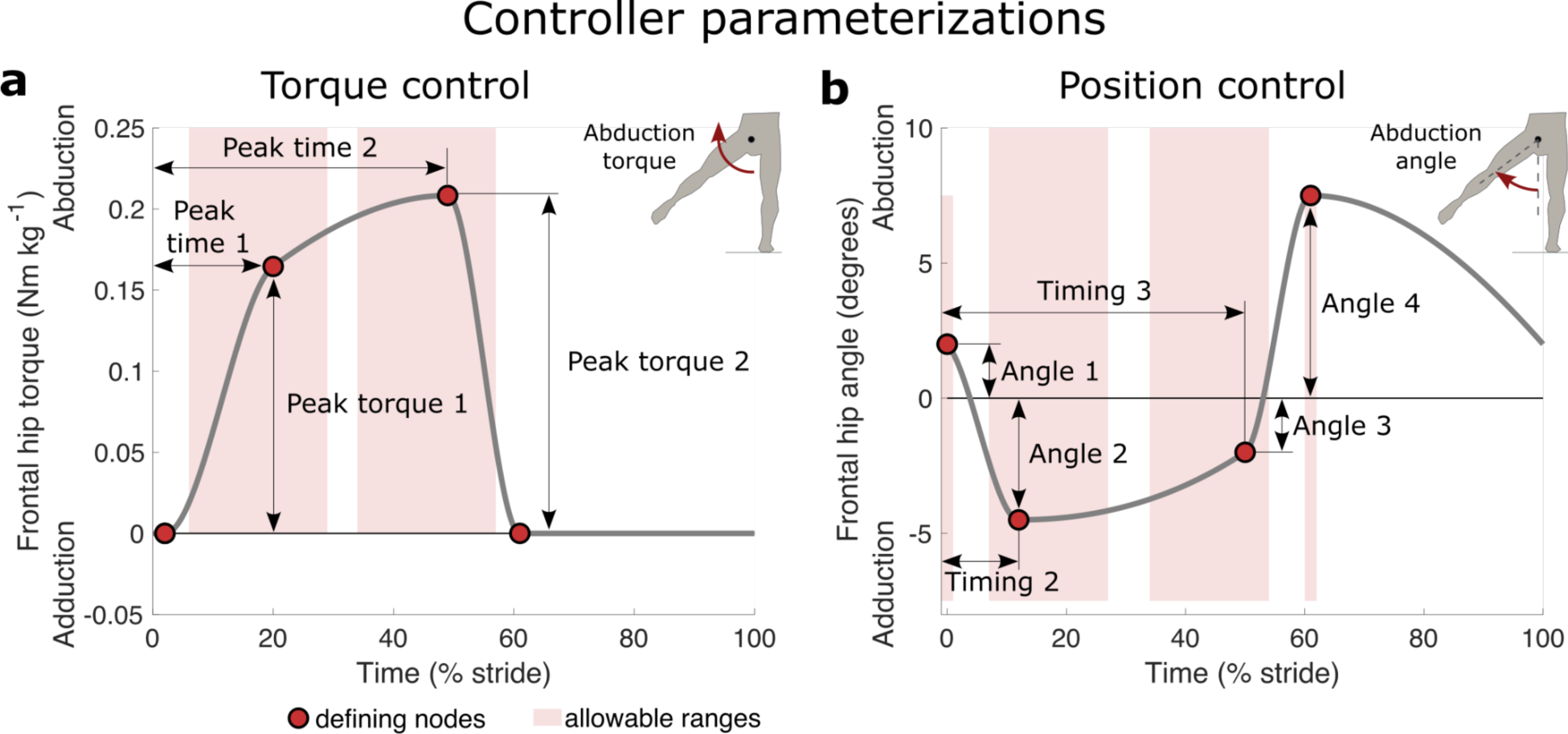
Controller parameterizations. **a,** Torque control parameterization. Hip abduction torque was commanded as a function of time, defined by a spline fit to four nodes (red circle). Four parameters (black arrow) were optimized during the experiment. **b,** Position control parameterization. Hip joint angle in the frontal plane was commanded as a function of time, defined by a spline fit to four nodes. Six parameters were optimized. Red shaded regions represent the allowable range of each parameter. 0% and 100% stride indicated the heel strike and the subsequent heel strike of the same leg, respectively.

To accurately track the desired torque, we implemented closed-loop feedback control using a proportional gain, a motor-velocity damping gain, a feedforward compensation gain for joint velocity, and a feedforward iterative learning gain, as detailed in [31]. The iterative learning process used errors from the corresponding point of preceding strides to anticipate the required corrections for the current stride [33]. The control signal derived from this feedback loop was then commanded to the motor drivers, operating in velocity control mode. This approach was accurate in applying torques, with root-mean-square (RMS) errors of 0.71 Nm (4.7 % of the maximum commanded torques).

When no torque was desired (i.e., walking in the exoskeleton without assistance or during the unassisted portion of the percent stride in the assisted condition shown in Fig. 2a), the motors tracked the position of the hip joint with a set amount of slack in the Bowden cable to prevent interference with the user. In this case, the RMS of the applied torque remained below 2.1 Nm, with its peak never exceeding 6 Nm in magnitude. Unintended torques were generated due to inertial effects and joint friction of the exoskeleton. These torque errors may also be present but unmeasured in many other exoskeletons, where sensors were typically placed in series with the transmission [19,20,34] rather than on the frame, as in our exoskeleton [31]. If we had employed such an approach, the torque during zero-torque mode would have been measured as negligible. During testing, we found the device comfortable for standing and walking in zero-torque mode.

#### Position Control

In the position control scheme, the exoskeleton was controlled by tracking frontal plane hip joint position trajectories as a function of the percent stride. This position profile was represented by a piecewise cubic Hermite interpolating polynomial spline with four nodes (Fig. 2b). The first and last nodes were fixed at heel strike and toe-off, respectively, as these specific timings have a significant impact on step width and circumduction. Two middle nodes were added to enable rapid changes in frontal plane hip angular velocity during the stance phase, as observed in unassisted walking [35]. These nodes also enable transitions between the lengthening and shortening of frontal hip muscles [36]. Consequently, six parameters with two constraints determined the magnitudes and timings of all four nodes, with minimum and maximum limits based on user comfort during pilot tests (Additional file 1: Table A2). To ensure trajectory continuity between strides, if the current stride time exceeded the average, the frontal hip angle at heel strike was enforced until the actual heel strike occurred. For shorter strides, the controller created a spline connecting the measured angle at heel strike with the desired angle at the second node.

To accurately track the desired position, we implemented closed-loop feedback control, similar to a method described in [37]. This involved calculating the commanded frontal hip angle as the sum of the desired angle, proportional feedback on angle error, and iteratively learned compensation [33]. The commanded frontal hip angle was then mapped from the joint encoder space to the abduction and adduction motor spaces using a ratio of the radius of sector drum pulley to the radius of the motor shaft. Subsequently, the control signal derived from this feedback loop was commanded to the motor drivers, operating in position control mode. This approach accurately tracked the prescribed position trajectories, resulting in RMS errors of 0.14 deg.

### Human-in-the-loop Optimization (HILO) Protocol

To optimize exoskeleton assistance for each control scheme, we used the Covariance Matrix Adaptation Evolutionary Strategy (CMA-ES) [38]. The objective of the optimizer was to minimize the metabolic cost of walking, and CMA-ES has shown success in reducing metabolic cost in previous exoskeleton studies [8,9,11,12,24]. During the experiment, a participant walked at 1.25m/s, and various patterns of exoskeleton assistance were provided by the CMA-ES algorithm. The algorithm sampled a generation of conditions from a distribution defined by parameter means and a covariance matrix, ranked the performance of the samples, and used these results to update the mean and covariance before sampling the next generation. In our setup, the performance of the samples referred to the metabolic cost of walking under the assistance provided in each particular condition. This metabolic cost was estimated using a first-order dynamics model of the human respiratory system after 2 minutes of walking [39].

In addition to the original CMA-ES algorithm, where all conditions were randomly sampled from a normal distribution, we introduced two additional conditions per generation. The first added condition was the parameter means for that generation, and the second added condition was the best-performing condition of the previous generation, following a similar approach to prior research [12]. This adjustment aimed to improve convergence by biasing the optimizer toward exploitation rather than exploration and also ensured that the optimizer didn’t deviate from any well-performing conditions. The remaining 8 or 9 conditions were sampled from a normal distribution, aligning with the core principles of the original CMA-ES algorithm. Consequently, the torque control scheme had 10 conditions per generation, and the position control scheme had 11 conditions per generation (Additional file 1: Table A3).

The duration of optimization was determined based on previous HILO studies, ensuring a balance between achieving convergence and maintaining experimental feasibility [8,11,12,30]. This experiment was conducted with an experienced user who is likely to adapt quickly to novel control schemes and new assistance strategies compared to novice users. For each optimization, the initial parameter means were set at the center of the parameter space to expedite convergence, irrespective of the ground truth optimum location (Additional file 1: Tables A1 and A2). Each control scheme underwent eight generations over two days (79 conditions for torque control and 87 conditions for position control), with each generation requiring 20 or 22 minutes of walking. The time gaps between optimization day 1 and day 2 were limited to within two days. To ensure algorithm convergence and avoid being trapped in a local minimum, we conducted HILO twice on the same participant for the torque control scheme and reported the result from both runs. The participant was allowed but not required to take breaks between generations. More details on the implementation of CMA-ES can be found in [8].

### Validation Protocol

To accurately assess the effect of the control scheme and assistance, a validation experiment was conducted on a separate day after each optimization. The intervals between optimization day 2 and the validation day were limited to five days to minimize potential time-related effects (e.g., optimal profile changes over time, training washout, etc.). During these experiments, we compared the performance of the optimized assistance with one or more of the following conditions: walking in the exoskeleton without assistance (No Torque), walking with assistance inspired by biological torque or angle (Bio), walking with simulation-inspired assistance (Sim), and walking without the exoskeleton (No Exo). The participant started by standing still for 4 minutes, followed by a 6-minute walk under each of the aforementioned conditions at a speed of 1.25 m/s. These conditions were then repeated in reverse order. To minimize donning and doffing time, we conducted the two No Exo conditions at the start and end of the protocol. The order of all other conditions was randomized. The participant was required to rest for at least three minutes between walking conditions to allow his metabolism to return to baseline.

#### Torque Control

We performed the optimization protocol twice under the torque control scheme, and we conducted a separate validation following each optimization run. After the first optimization run, we evaluated two conditions: Optimized Torque and No Torque. After the second optimization run, we evaluated five conditions: Optimized Torque, No Torque, No Exo, Bio Torque, and Sim Torque. The Bio Torque condition delivered a profile that mimicked a scaled-down version of the biological hip moment in the frontal plane, using reference data [5]. The Sim Torque condition delivered a profile inspired by the simulation study by Dembia et al. [6].

#### Position Control

Following the optimization protocol under the position control scheme, we evaluated four conditions: Optimized Angle, No Torque, No Exo, and Bio Angle. The Bio Angle condition delivered a profile that mimicked the biological hip angle in the frontal plane, based on reference data [35].

### Measured Outcomes and Analysis

During validation, we measured metabolic cost, applied torque, exoskeleton joint angles, and ground reaction forces and moments for each walking condition. These measurements were then averaged over the final three to six minutes of each condition to ensure that the user’s metabolic rate and gait had reached a steady state. All conditions were evaluated twice, and the measurements were averaged across both evaluations. Since all experiments were conducted on a single subject, differences between conditions were analyzed without formal statistical tests.

#### Metabolic Cost

We calculated metabolic cost using indirect calorimetry (Quark CPET, Cosmed, Italy). We measured oxygen consumption, carbon dioxide production, and breath duration on a breath-by-breath basis. Metabolic rate was calculated for each condition using the Brockway equation [40], along with a weighted average based on breath duration. To calculate the metabolic cost of walking, we subtracted the cost of quiet standing from the walking conditions. To ensure accurate metabolic measurements, the participant fasted for four hours prior to the testing.

#### Torques and Kinematics

Exoskeleton torques and joint angles were measured using strain gauges and absolute encoders [31]. To remove the joint angle offset, the encoder readings taken during the natural standing position were subtracted from the subsequent readings. These angles were used to estimate user kinematics, assuming minimal relative movement between the user and the exoskeleton. The measurements were averaged across both legs.

## Results

### Torque Control

An optimized torque profile from HILO did not reduce the metabolic cost of walking. Both the first and second peak torque magnitudes decreased significantly, approaching zero (Additional file 1: Figs. A1a and A1b), resulting in the optimized profile similar to the No Torque condition (Fig. 3a). Consequently, the metabolic cost difference between the No Torque and Optimized Torque conditions was only 0.8% (No Torque: 4.31 W kg^-1^; Optimized Torque: 4.35 W kg^-1^; Fig. 3b), which fell within the range of measurement noise [41]. To ensure algorithm convergence and prevent getting trapped in a local minimum, we conducted HILO twice on the same participant. In the second run, peak torques once again decreased toward zero, consistent with the results of the first run (Fig. 3a). Although the peak timings did not converge to the same value in both runs (Additional file 1: Figs. A1c and A1d), the near-zero peak torque magnitudes meant that peak timings had no impact on the assistance profile (Additional file 1: Table A1). The bio-inspired and simulation-inspired assistance substantially increased the metabolic cost by 13.4% and 27.1%, respectively, compared to the No Torque condition (Bio. Torque: 4.75 W kg^-1^; Sim. Torque: 5.32 W kg^-1^; Fig. 3c). Walking without the assistance (No Torque) resulted in a 40.0% increase in net metabolic cost relative to walking without the exoskeleton (No Exo: 2.99 W kg^-1^; Fig. 3c).

**Fig. 3.**
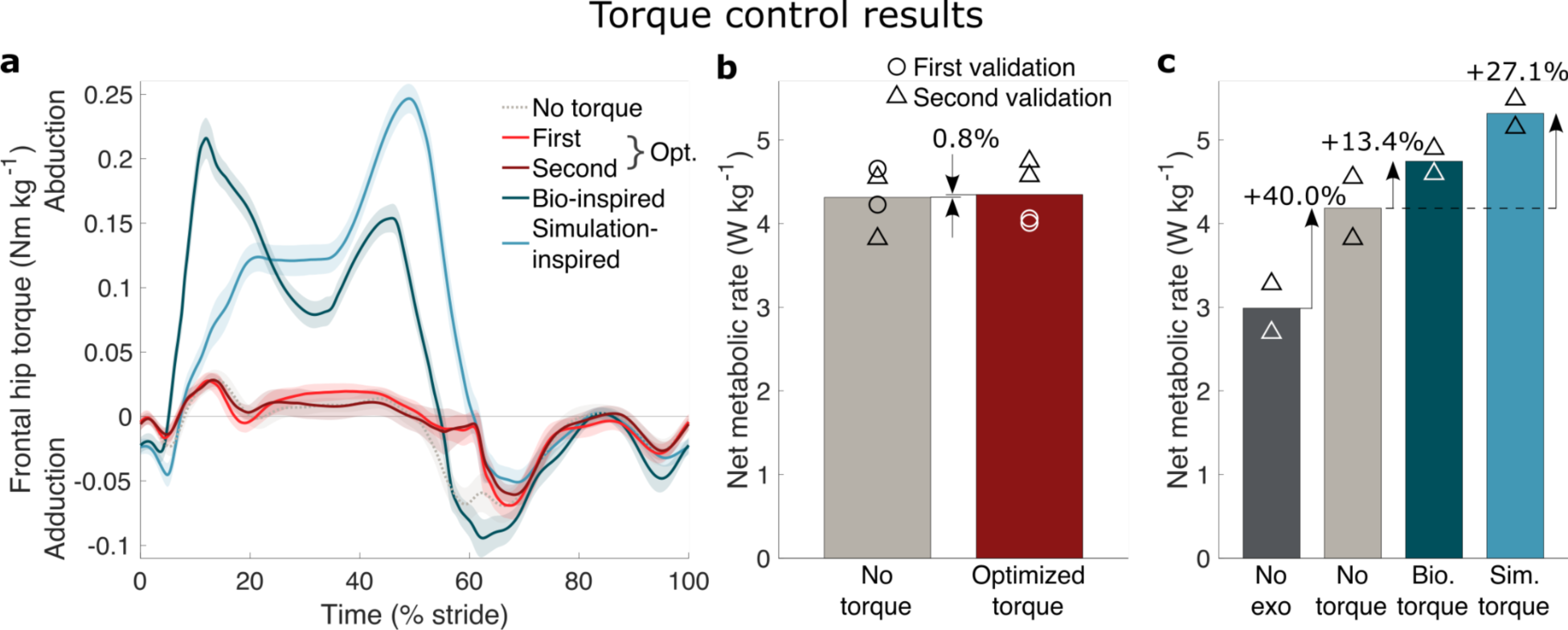
Torque control results. **a,** Measured exoskeleton torques at the hip joint in the frontal plane. Lines and shaded regions represent the mean and standard deviations, respectively, calculated from both legs and across the duration of all corresponding walking trials in the validation protocol. **b,** Metabolic results for the optimized torque (red) and no torque (light gray) conditions. Each walking bout in the first and second validations is indicated by the symbols. **c,** Metabolic results for other conditions (no exoskeleton: dark gray, no torque: light gray, biological torque: teal, simulation torque: sky-blue).

### Position Control

An optimized position profile from HILO also did not reduce the metabolic cost of walking. The profile was nearly constant, varying within 2 degrees around the neutral hip angle (0°) in the frontal plane throughout the gait cycle (Fig. 4a). Interestingly, despite differences in frontal hip angle and torque between the Optimized Angle condition and the No Torque condition (Fig. 4a and Additional file 1: Fig. A2), both conditions exhibited very similar metabolic rates. The difference in metabolic cost between the No Torque and Optimized Angle conditions was only 2.8% (No Torque: 3.58 W kg^-1^; Optimized Angle: 3.68 W kg^-1^; Fig. 4b), which fell below the minimal detectable change threshold [42]. A position profile that mimicked the biological hip angle in the frontal plane increased the metabolic cost by 17.8% compared to the No Torque condition (Bio. Angle: 4.22 W kg^-1^; Fig. 4c). Walking without any assistance (No Torque) resulted in a 22.3% increase in net metabolic cost compared to walking without the exoskeleton (No Exo: 2.93 W kg^-1^; Fig. 4c).

**Fig. 4.**
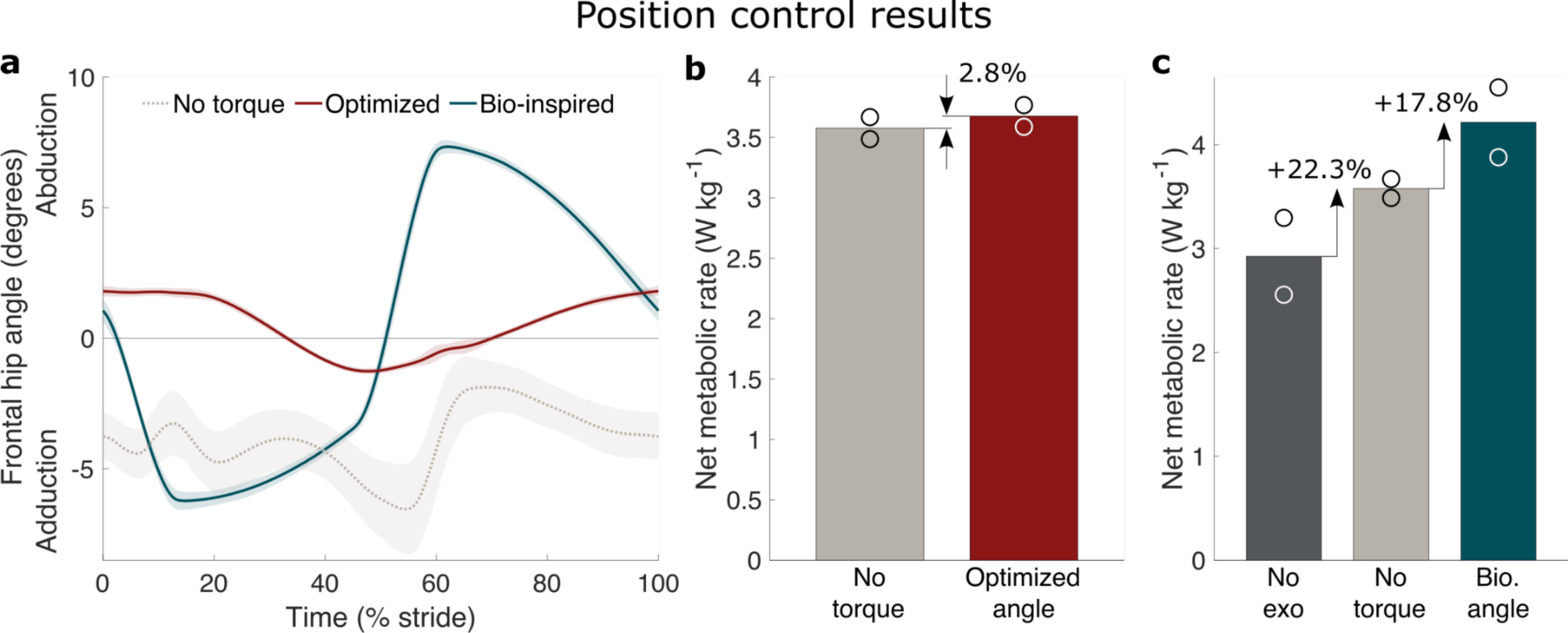
Position control results. **a,** Measured exoskeleton angle at the hip joint in the frontal plane. Lines and shaded regions represent the mean and standard deviations, respectively, calculated from both legs and across the duration of all corresponding walking trials in the validation protocol. **b,** Metabolic results for the optimized angle (red) and no torque (light gray) conditions. Each walking bout is indicated by the symbols. **c,** Metabolic results for other conditions (no exoskeleton: dark gray, no torque: light gray, biological angle: teal).

## Discussion

None of the frontal-plane hip assistance approaches reduced the metabolic cost of walking in this human-in-the-loop optimization study. The optimal torque magnitude was identified to be near zero, indicating that any hip abduction torque would increase metabolic rate. The optimized position profile remained nearly constant throughout the gait cycle, resulting in no metabolic reduction. Although our study did not encompass an exhaustive exploration of all possible control approaches, reducing the metabolic cost through frontal hip assistance does not seem promising.

The lack of metabolic reduction does not appear to be due to a malfunction of the optimization algorithm or insufficient exoskeleton assistance training. Algorithm convergence was ensured by running the CMA-ES algorithm for a similar number of generations/duration as other successful HILO studies (e.g., 4–12 generations to optimize 3–8 parameters at the hip [11,12] or ankle [8,12] in the sagittal plane). Additionally, repeating HILO on the same participant consistently converged to the same optimized profile, indicating the algorithm’s reliability and consistency. Further, the optimized profile demonstrated a substantial improvement over the bio-inspired and simulation-inspired profiles, which are prominent in the field. This improvement was observed in both torque and position control schemes. Lastly, the participant was an experienced exoskeleton user, with tens of hours of exposure to the exoskeleton assistance used in this study; therefore, the lack of training does not explain the failure of metabolic reduction. Overall, this evidence collectively implies that a 0% reduction may be the best achievable performance with frontal hip assistance, demonstrating that reducing the metabolic cost of walking through frontal-plane hip assistance may not be possible.

This study provides empirical support indicating the greater challenge of assisting tasks that do not require substantial positive power. Concentric muscle contractions are known to be four times more metabolically expensive than eccentric contractions [43], and there is a strong correlation between metabolic reduction and positive work performed by exoskeletons [8,17]. However, due to the limited range of frontal hip motion, the positive work over one stride of hip abduction/adduction is only about 0.09 J/kg, which is five times smaller than that of the hip (0.48 J/kg) or ankle (0.42 J/kg) in the sagittal plane [5]. Achieving metabolic reduction for tasks that require large amounts of positive power might be easier, because of the potential for the exoskeleton to replace expensive positive power production by muscles [44–47]. (It should be noted, however, that more power is not always better [7,8,11,48].) On the other hand, assisting tasks that don’t require much positive power (e.g., maintaining balance) could be more difficult due to the task’s reliance on neural feedback control, involving factors such as body state information, prediction of potential perturbations, and neuromuscular coordination in response to disturbances. Exoskeletons that lack access to this crucial information may not be able to effectively coordinate with humans in this type of task.

Our experimental results are inconsistent with the musculoskeletal simulation, which predicted a 13% metabolic reduction through hip abduction assistance [6]. This discrepancy can be attributed, in part, to differences in the tasks and assumptions made during the simulation. The simulation was conducted under heavily loaded walking conditions (38 kg load on the torso), whereas we tested unloaded, normal walking in the experiment. Additionally, the simulation lacks an explicit model of the human nervous system, potentially resulting in an inadequate representation of human adaptation. For instance, it assumed fixed kinematics (and ground reaction forces), but our observations revealed kinematic changes associated with assistance (Additional file 1: Fig. A3). The simulation’s prediction of muscle activity during exoskeleton assistance can also be inaccurate [49]. Humans may behave differently from the simulation’s approach of minimizing the sum of squared muscle activations, possibly due to factors such as fatigue, joint pain, and/or neural coordination [50]. To enhance prediction accuracy, the musculoskeletal simulation could integrate factors like these into its modeling.

The metabolic difference between the No Torque and No Exo conditions was substantial. On average, over two different validation days in torque and position control, the No Torque condition increased metabolic rate by 31.3% compared to No Exo (No Exo: 2.96 W kg^-1^; No Torque: 3.88 W kg^-1^). This increase can be attributed to the metabolic penalty of carrying the additional mass of the exoskeleton, which could account for an estimated 23.1% increase in metabolic rate (0.68 W kg^-1^) [51]. Furthermore, the absence of arm swing during the No Torque condition can contribute to an approximate additional 9.4% increase (0.28 W kg^-1^) [52]. Other factors, such as circumduction [53] and increased step width and its variability [54], may also contribute to the observed differences.

We investigated the feasibility of frontal hip assistance in two different control schemes with repeated measurements, but all of them were tested on a single subject. Given the large inter-subject variability observed in other exoskeleton studies, we refrain from making strong claims about its generalizability. However, it is worth noting that exoskeletons demonstrating large metabolic reduction showed consistent reductions at the individual level [7–9,13]. Furthermore, the subject in our study was an expert exoskeleton user, who also participated in a variety of other sagittal hip, knee, and ankle exoskeleton/exosuit studies [10,11,13,23,29,45,46,55–58] and achieved an average metabolic reduction among all participants. Thus, increasing the number of subjects is unlikely to change the study’s conclusion.

## Conclusions

This study optimized frontal-plane hip assistance through human-in-the-loop optimization in torque and position control schemes, but found no reduction in the metabolic cost of walking. The disparity between our experimental results and a prior simulation [6] underscores potential opportunities to improve prediction accuracy in musculoskeletal simulations. Future experimental work should investigate hip abduction assistance during loaded walking. Alternative control strategies and alternative objectives should also be explored, for example a neuromuscular controller optimized to improve walking balance. With such approaches, it may yet be possible for an exoskeleton with hip abduction assistance to reduce the effort required for walking.

## List of abbreviations

HILO: Human-in-the-loop optimization
GRF: Ground reaction forces
CoP: Center of pressure
RMS: Root-mean-square
CMA-ES: Covariance Matrix Adaptation-Evolutionary Strategy
Bio: Biological
Sim: Simulation-inspired
No Exo: No Exoskeleton
Opt: Optimized

## Supplementary Information

**Additional file 1.** This file provides additional data not shown in figures and tables in the main text. **Figure A1** shows optimized parameters across HILO generations in the torque control scheme. **Figure A2** shows frontal hip torque of the exoskeleton in the position control scheme. **Figure A3** shows frontal hip angle of the exoskeleton in the torque control scheme. **Table A1** provides HILO parameter ranges, initial seeds, and optimized values in the torque control scheme. **Table A2** provides HILO parameter ranges, initial seeds, and optimized values in the position control scheme. **Table A3** provides overview of the protocol for each human-in-the-loop optimization (HILO) case study.

## Declarations

### Ethics approval and consent to participate

All experiments were approved by the Institutional Review Board at Stanford University (Protocol #: IRB-57846). The participant provided written and informed consent to participate.

### Consent for publication

The participant provided written and informed consent for the publication of the data and images.

### Availability of data and materials

All data generated or analyzed during this study are included in this published article.

### Authors’ contributions

J.K. and S.H.C. conceived of the study concept and experimental methods. J.K. developed the exoskeleton controller and conducted the human subject experiments. J.K. analyzed the data and prepared the manuscript. M.R., C.K.L., and S.H.C. edited and provided feedback on the manuscript. All authors read and approved the final manuscript.

## Acknowledgments

The authors would like to thank Atharva Wadhokar for his valuable assistance in data collection, as well as the entire Stanford Biomechatronics Laboratory for their insightful feedback and suggestions.

