## Additional file 1 for "Frontal hip exoskeleton assistance does not appear promising for reducing the metabolic cost of walking: A preliminary experimental study"

### Additional file 1 – Supplementary Figures & Tables

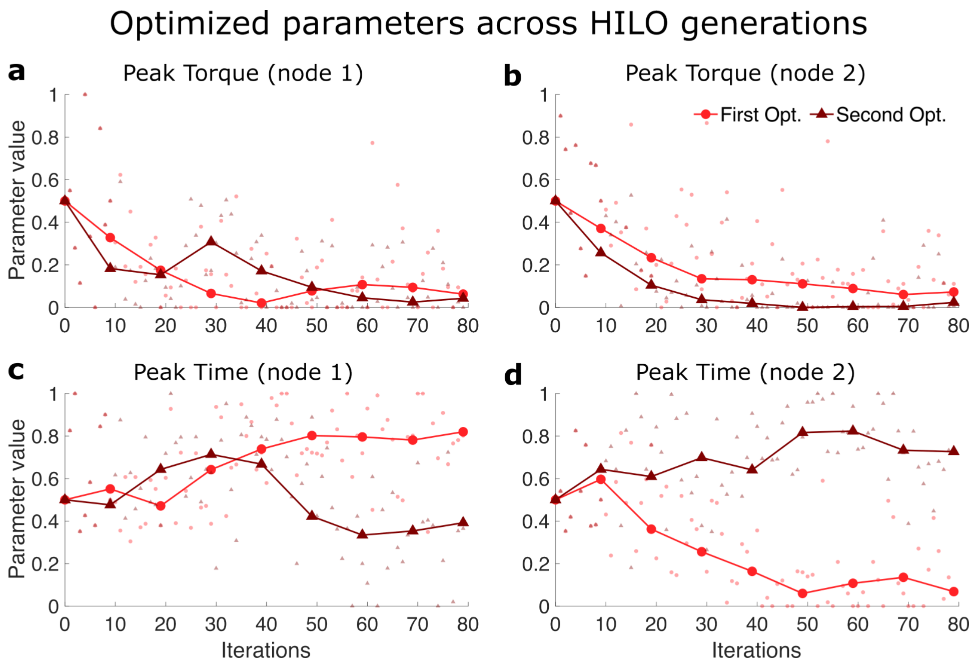


**Fig. A1.** Optimized parameters across HILO generations in the torque control scheme. **a,** The magnitude of the first peak node. **b,** The magnitude of the second peak node. **c,** The timing of the first peak node. **d,** The timing of the second peak node. Each parameter value is normalized such that 0 and 1 represent the minimum and maximum values within the allowable range. Each optimization run is indicated by the symbols. Large markers with lines indicate the parameter means of each generation, whereas small markers indicate the parameter values at each iteration.

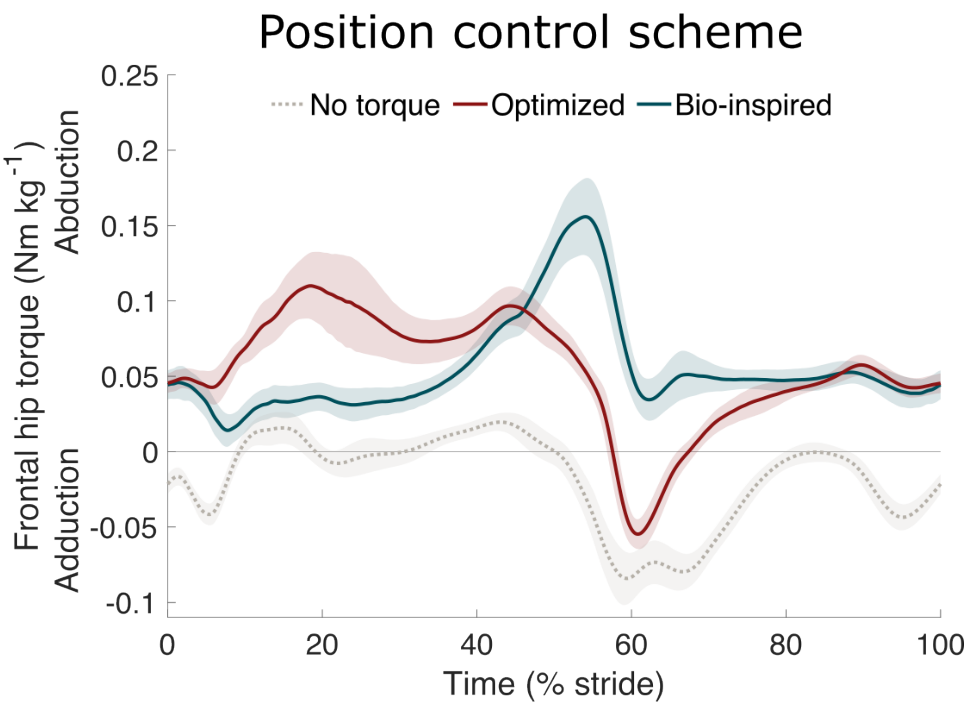


**Fig. A2.** Frontal hip torque of the exoskeleton in the position control scheme. Lines and shaded regions represent the mean and standard deviations, respectively, calculated from both legs and across the duration of all corresponding walking trials in the validation protocol.

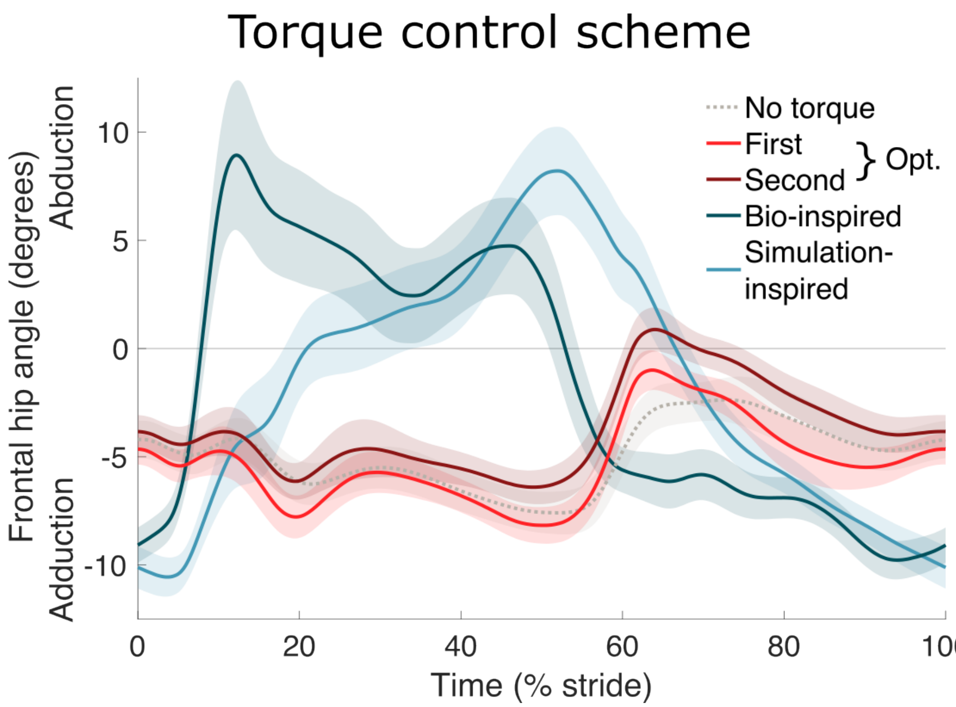


**Fig. A3.** Frontal hip angle of the exoskeleton in the torque control scheme. Lines and shaded regions represent the mean and standard deviations, respectively, calculated from both legs and across the duration of all corresponding walking trials in the validation protocol.

**Table A1**. HILO Parameter ranges, initial seeds, and optimized values in the torque control scheme.

| | Peak torque 1 (Nm $kg^{-1}$) | Peak torque 2 (Nm $kg^{-1}$) | Peak time 1 (% stride) | Peak time 2 (% stride) |
|---|---|---|---|---|
| Minimum | 0 | 0 | 6.0 | 34.0 |
| Initial seed | 0.13 | 0.13 | 17.5 | 45.5 |
| Maximum | 0.25 | 0.25 | 29.0 | 57.0 |
| Optimized ($1^{st}$) | 0.02 | 0.02 | 24.9 | 35.6 |
| Optimized ($2^{nd}$) | 0.01 | 0.01 | 15.0 | 50.7 |

**Table A2**. HILO Parameter ranges, initial seeds, and optimized values in the position control scheme.

| | Angle 1 (deg) | Angle 2 (deg) | Angle 3 (deg) | Angle 4 (deg) | Timing 2 (% stride) | Timing 3 (% stride) |
|---|---|---|---|---|---|---|
| Minimum | -7.5 | -7.5 | -7.5 | -7.5 | 7.0 | 34.0 |
| Initial seed | 0 | 1.3 | 1.3 | 0 | 17.0 | 44.0 |
| Maximum | 7.5 | 10.0 | 10.0 | 7.5 | 27.0 | 54.0 |
| Optimized | 1.9 | 1.9 | -1.1 | -0.5 | 15.3 | 47.6 |

**Table A3**. Overview of the protocol for each human-in-the-loop optimization (HILO) case study. It includes the control scheme, the number of parameters (params) in the control architecture, the number of control laws per generation (gen), the number of generations, the number of participants, and the number of HILO runs. Two additional control laws per generation (i.e., +2) represent the parameter means for that generation and the best-performing condition from the previous generation.

| Control scheme | Params (#) | Control laws/gen (#) | Gens (#) | Participants (#) | Runs (#) |
|---|---|---|---|---|---|
| Torque | 4 | 8+2 | 8 | 1 | 2 |
| Position | 6 | 9+2 | 8 | 1 | 1 |
